# Analysis of single-cell RNA-sequencing data to identify quiescent and proliferating neural cell populations in Glioblastoma

**DOI:** 10.1101/2021.12.09.472030

**Authors:** Rajeev Vikram, Wen□Cheng Chou, Pei-Ei Wu, Wei-Ting Chen, Chen-Yang Shen

## Abstract

**Background:** Diffuse Glioblastoma (GBM) has high mortality and remains one of the most challenging type of cancer to treat. Identifying and characterizing the cells populations driving tumor growth and therapy resistance has been particularly difficult owing to marked inter and intra tumoral heterogeneity observed in these tumors. These tumorigenic populations contain long lived cells associated with latency, immune evasion and metastasis.

**Methods:** Here, we analyzed the single-cell RNA-sequencing data of high grade glioblastomas from four different studies using integrated analysis of gene expression patterns, cell cycle stages and copy number variation to identify gene expression signatures associated with quiescent and cycling neuronal tumorigenic cells.

**Results:** The results show that while cycling and quiescent cells are present in GBM of all age groups, they exist in a much larger proportion in pediatric glioblastomas. These cells show similarities in their expression patterns of a number of pluripotency and proliferation related genes. Upon unbiased clustering, these cells explicitly clustered on their cell cycle stage. Quiescent cells in both the groups specifically overexpressed a number of genes for ribosomal protein, while the cycling cells were enriched in the expression of high-mobility group and heterogeneous nuclear ribonucleoprotein group genes. A number of well-known markers of quiescence and proliferation in neurogenesis showed preferential expression in the quiescent and cycling populations identified in our analysis. Through our analysis, we identify ribosomal proteins as key constituents of quiescence in glioblastoma stem cells.

**Conclusions:** This study identifies gene signatures common to adult and pediatric glioblastoma quiescent and cycling stem cell niches. Further research elucidating their role in controlling quiescence and proliferation in tumorigenic cells in high grade glioblastoma will open avenues in more effective treatment strategies for glioblastoma patients.

## INTRODUCTION

Diffuse gliomas are tumors of the central nervous system with histological similarity to glial cells. Worldwide, approximately 100,000 new cases of diffuse glioma are reported every year[1]. Despite it being a relatively rare cancer type, diffuse gliomas have a very poor prognosis with high mortality burden. The 2016 WHO classification of gliomas divides them into astrocytomas, oligodendrogliomas and oligoastrocytomas with subgroupings based on IDH mutations and 1p19q co-deletion status[2]. Irrespective of the categories, the tumors are graded from one to four according to the histological degree of malignancy.

The grade IV diffuse astrocytoma (IDH-wildtype) also called as glioblastoma (GBM) accounts for about 75% of all diffuse gliomas with a median survival of about one to two years after therapy, making it the most lethal of gliomas[2, 3]. Intratumoral heterogeneity in GBM is a key challenge to developing effective therapeutic strategies. Neurodevelopmental bi-lineage hierarchy does partially explain the heterogeneity in IDH-mutant and pediatric gliomas, however, this bi-lineage hierarchy model fails to explain the widespread phenotypic heterogeneity and evolving phenotypic states in GBM. The cancer stem cells (CSC) theory suggests that Glioblastoma Stem-like Cells (GSCs) are at the center of the tumor organization and instrumental in generating and replicating intratumoral phenotypic heterogeneity[4]. Indeed, similar to other cancers, intratumoral heterogeneity, resistance to treatment and relapse in GBM has been attribute to this small subpopulation in a number of studies. Although GSCs seem to be a key target in GBM therapy, their existence and cellular nature remains a hotly debated topic. While there is evidence pointing to the presence of such GSCs, identification of these cells has remained a challenge. Primarily because there is no marker which can be considered to universally identify GSCs[5].

In most cancers including GBM, single surface marker approach has been used to identify CGSCs. A number of cell membrane antigens like CD133, CD15/SSEA, CD44, PDGFRA, EGFR or A2B5 are shown to be associated with potential GSCs. Earlier studies showing the tumorigenic potential of cells isolated using one or a combination of these markers [6–11] did not address the tumorigenic potential of marker negative cells. Later studies however show that both the marker positive and negative glioma fractions can show multipotent behavior [8, 12]. Recent reports show that marker negative cells are able to generate marker positive cells and replicate the tumor heterogeneity[13]. Thus the evidence so far indicates a non-hierarchical model where cells niches with strong cellular plasticity are at the core of recreating intratumoral heterogeneity. Evidence supporting intratumoral cell niches with high cellular plasticity in glioblastoma comes from a recent study by Jung, Erik, et al.[14]. The authors show the existence two complementary cellular niches driving tumor progression and therapy resistance in GBM. Further evidence of the plastic nature of GSC niches come from studies which demonstrate the generation of potential GSCs from non-tumorigenic glioma cells [15]. Based on these and a number of other recent studies, it seems that within the tumor, potential GSCs remain in interactive niches and are highly plastic in addition to being able to acquire therapy resistance and tumorigenicity. Recent studies in tumor immunosurveillance evasion suggests that these niches are composed of small populations of cycling and quiescent stem cell like cells [16].

Advances in newer methods to study cellular transcriptomes at the single cell level especially massively parallel single-cell RNA-sequencing (scRNAseq) has significantly enhanced our understanding of spatial and temporal heterogeneity in glioblastoma. Recent single cell studies of gliomas have shown the presence of a progenitor type population [17–19]. Interestingly, in terms of conventional markers, tumors from different patients show variability in their expression, suggesting heterogeneity within GSCs[20]. As more and more genomic and transcriptomic data from single cell experiments in gliomas becomes available, it is becoming more evident that although canonical GSC markers seem to be associated with proliferative cells in low grade gliomas, such correlation is not evident in GBM[21]. Thus, projects designed to identify GSCs on the hierarchical CGSC model have largely been ineffective.

To develop a better diagnostic and treatment strategy for GBM as well as low grade glioma, is it important understand the dynamics of the tumor microenvironment, especially the intrinsic plasticity of the cell niches. Understanding the mechanism of maintenance of these highly plastic cell subpopulations within the tumor, the role of the microenvironment dynamics in selection and survival of such populations and their propagation are instrumental in unearthing the reasons for the development of resistance to Temozolomide (TMZ) chemotherapy and radiotherapy.

A number of recent studies have utilized scRNAseq to study gliomas of different origin and grade generating a wealth of data on the transcriptomic nature of cells within the tumor[14, 17, 19, 22]. While these studies primarily focused on different states of gliomas and tumor-immune cell interaction, few studies have tried to delineate differences in cellular states within neural cells in GBM. Reanalysis of these scRNA-seq datasets can give us a deeper understanding of the genomic and transcriptomic commonalities within malignant neural subpopulations. Identification of GSC niches with quiescent population perhaps holds and the key to developing strategies which can target genes/transcripts involved in maintaining GSC plasticity and crosstalk in high grade gliomas.

Here, we analyze the scRNAseq data from four different studies encompassing thousands of tumor and peripheral cells from pediatric and adult IDH-wildtype glioblastoma patients to identify and study quiescent and cycling GSCs. The tumor types range from primary to relapsed tumors. Our results show that cycling and quiescent like-cell subpopulation are present in most GBM tumors with a gene expression signature associated with ribosomal biogenesis, cell cycle activation and malignancy. Key overexpressed genes include *DCX, SOX4* and *DLL3*, known markers for quiescence, stemness and tumorigenicity. These cells have a Copy number variance pattern distinguishing them from other neural cells subtypes within the tumor. and key genes/transcripts expression pattern in these niches across GBM tumor types.

## RESULTS

### Selection of datasets and identification of neural cells

To identify common GSC like populations across GBMs, we selected scRNAseq datasets from different studies representing IDH-wildtype, grade IV glioblastomas. Included datasets represent major patient groups (pediatric, adult and recurrent). Table 1 shows the major characteristics of the included datasets. For differential gene expression analysis of GBM subpopulations, we also included brain metastasis (lung squamous cell carcinoma) data set from the study GSE117891.

**Table 1.**
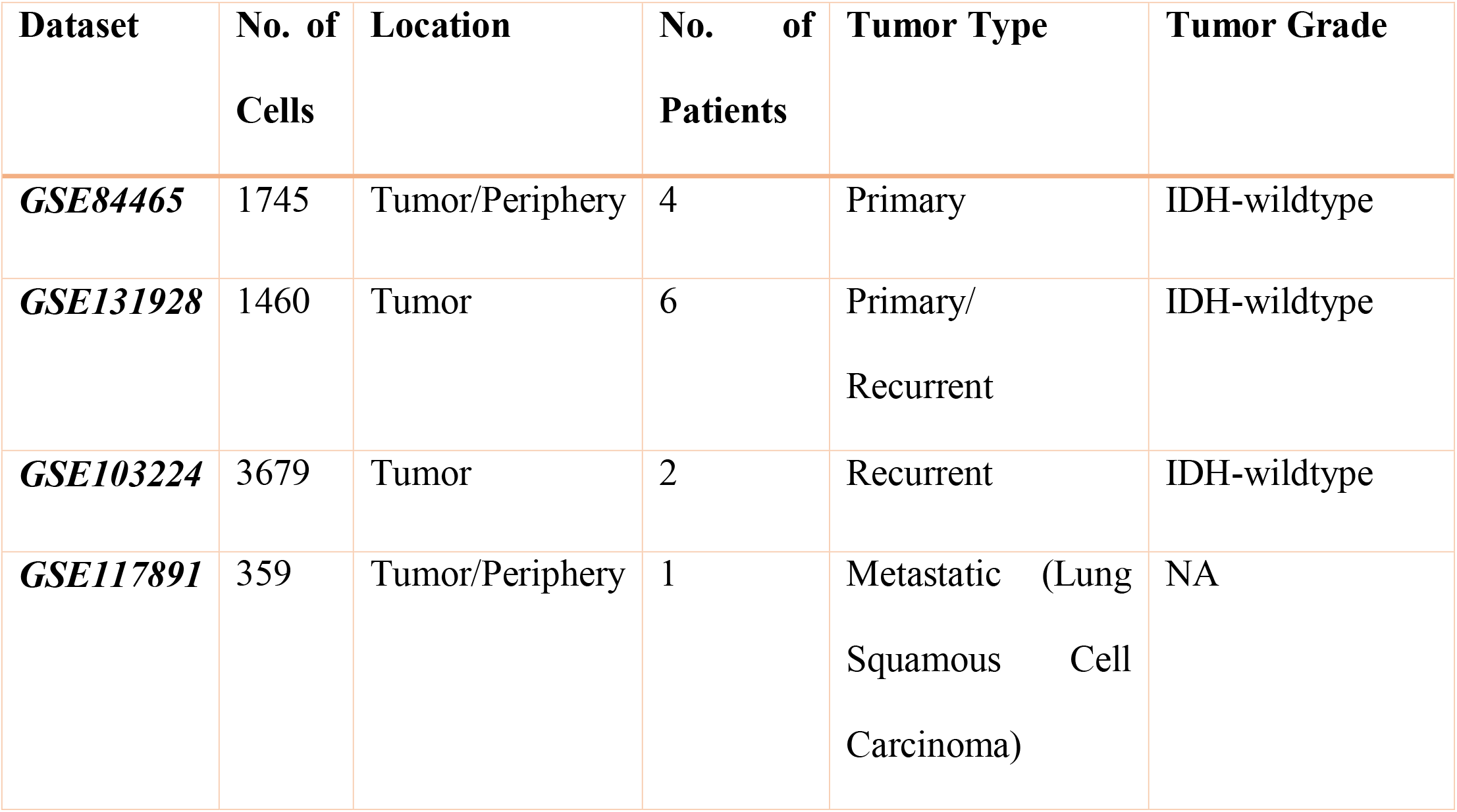
Dataset Summary.

| Dataset | No. of Cells | Location | No. of Patients | Tumor Type | Tumor Grade |
| --- | --- | --- | --- | --- | --- |
| ***GSE84465*** | 1745 | Tumor/Periphery | 4 | Primary | IDH-wildtype |
| ***GSE131928*** | 1460 | Tumor | 6 | Primary/<br>Recurrent | IDH-wildtype |
| ***GSE103224*** | 3679 | Tumor | 2 | Recurrent | IDH-wildtype |
| ***GSE117891*** | 359 | Tumor/Periphery | 1 | Metastatic (Lung<br>Squamous Cell<br>Carcinoma) | NA |

Significant inter and intratumoral heterogeneity is a challenge in identifying GSC like niches because gene expression patterns of different cell types and sample origin (i.e., transcriptomic diversity of the samples) induce strong variance which often masks the similarities in cellular programs in small subpopulations. Malignant cells with stem cell like properties are typically a very small subset of the tumor population. We therefore first sought to enrich our datasets based on clear cell identity and malignancy. The datasets were first clustered using the standard Seurat-sctarnsform pipeline. As, GBMs primarily originate in neural cells, we first identified the clusters of clear neural/glial (neuronal) or myeloid/immune origin by measuring the expression of known markers for immune cells (*CD4, CD83 & HLA-DRA*) and neuronal cells (*S100B, OLIG1 & SCG2*) [23, 24] (Figure 1 B, C). Clusters are represented using Uniform Manifold Approximation and Projection (UMAP) (Figure 1A). The neuronal cluster significantly overexpressed *EGFR*, a gene overexpressed in 30-50% of all GBMs and associated with neoplasia. Interestingly, non-GBM tumor cells (GSE117891) did not express EGFR and showed markedly different non myeloid/immune cluster profile. (Figure 1D). These clusters showed distinct expression profiles for glial cell markers for astrocyte(S100B) and oligodendrocytes(OLIG2). Indicating the presence of transformed non neural cells in these clusters (Figure 1 B, C).

**Figure 1:**
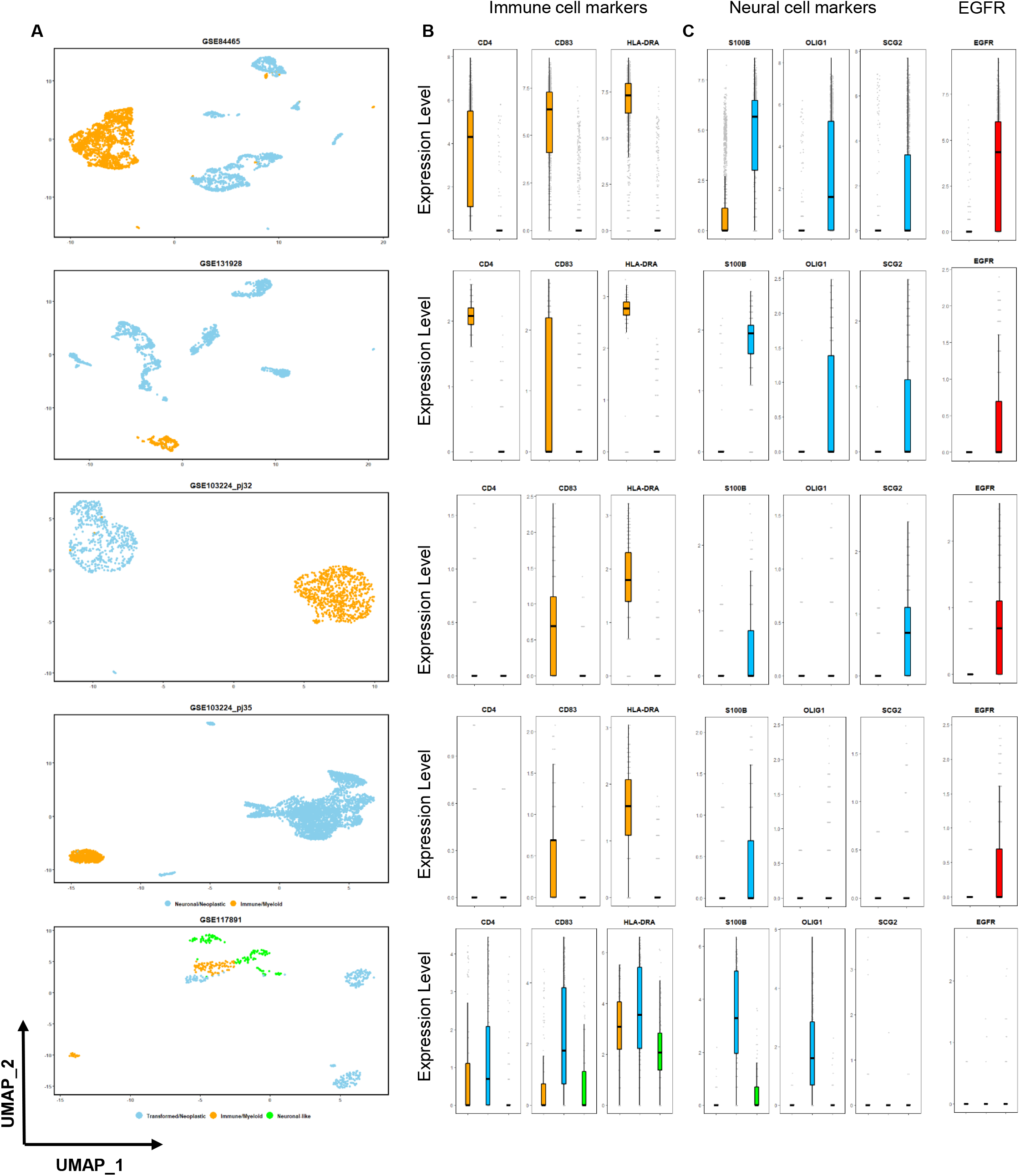
Identification of Immune and neural cells clusters (A) UMAP representation of clusters of all datasets. Immune cells are colored yellow; neural cells are colored blue, for GSE117891, blue represents transformed cluster, yellow-immune and green, neuronal-like. (B) Bar plot showing cluster wise expression levels of immune cell markers, bars are color coded according to the clusters in A. (C) Bar plot showing expression levels of neuronal markers, bars are color coded according to clusters in A.(D) Bar plot showing expression levels of EGFR, bars are colored red. All expression values are log transformed.

### Integrated cluster analysis

To identify malignant or transformed stem-like cells, we decided to focus exclusively on neuronal cells and examine them in detail. To do so, we first removed the immune/myeloid clusters from the study. Next, we used the mutual nearest neighbors (MNNs) method in conjugation with Seurat to integrate the datasets. For comparative analysis, we created two integrated datasets comprising of adult and pediatric GBM patients. Table 2 shows the patient wise detail of the adult and pediatric groups. Both adult and pediatric groups clustered largely according to cell types and cell stages (figure 2 A, B). Comparison of adult and pediatric groups showed that a number of genes specific to cell cycle, neuronal and glial cells like *TOP2A, SLC1A2, SSR4, APOD, PLP1, CD74, RTN1, HMGB2, CHI3L1, SERPINE1* were commonly overexpressed in certain clusters in both the groups. (figure 2 B). Whilst SLC1A2, APOD, PLP1, CD74, and RTN1 are cell type specific proteins and expressed by astrocytes, oligodendrocytes, OPCs and endothelial cells respectively[25], TOP2A and HMGB2 expression is specific to transcriptional activation and cell cycle[26, 27]. CHI3L1 and SERPINE1 are both proteins related to cell differentiation and malignant transformation[28, 29]. Comparatively, Adult GBM dataset exhibited greater number of distinct clusters which might be reflective of higher cell type and disease stage diversity. The presence of specific clusters overexpressing TOP2A and SOX4 in both adult and pediatric groups indicated a similarity in gene expression profile of these clusters across groups. Cycling or actively dividing and quiescent stem cells are known to coexist in adult stem cell niches[30]. Hence, we assessed the expression distribution of known markers for proliferation (KI67and CD44) and maintenance of cellular plasticity (SOX11 and DCX). The expression of these genes coincided with clusters1,2 and 3 the adult group while in the pediatric group, all clusters except clusters 3 and 7 showed high expression of these genes (Figure 2 C). Interestingly, CD44 expression did not follow this pattern, its expression was confined to clusters unrelated to the expression of other markers. Indeed, recent reports have questioned the validity of CD44 as a reliable marker of proliferation in glioblastoma [20, 31]. As, *TOP2A, MKI67, DCX,SOX4* and *SOX11* genes are known to show high expression in cycling and Quiescent stem cells respectively[26, 32], we suspected the presence of similar niches within the above mentioned clusters in the GBM groups. To verify this hypothesis, we decided to further analyze the cell type context of these clusters. Top 50 differentially expressed genes for both adult and pediatric groups are included in additional file 1.

**Table 2.**
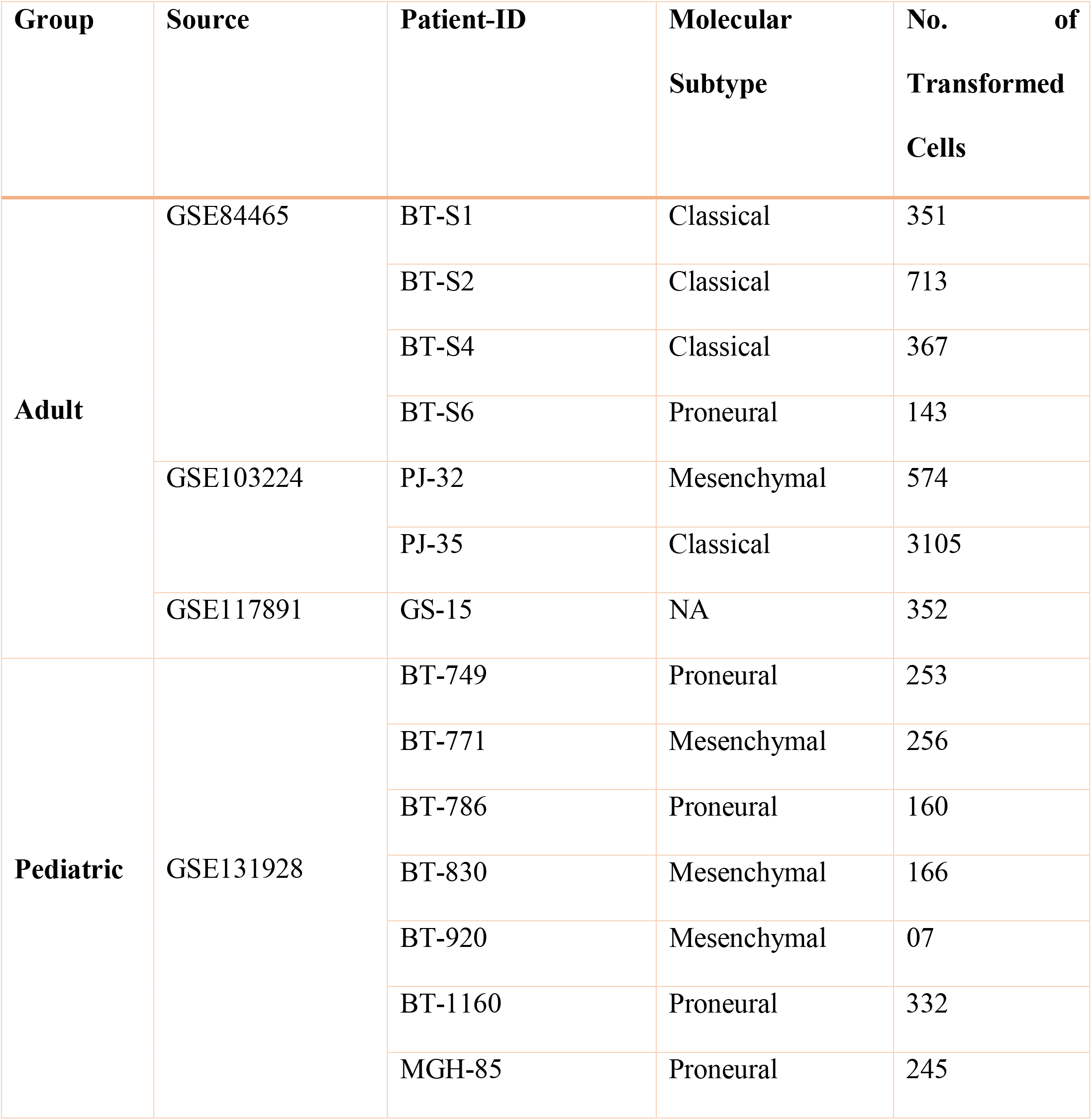
Study Group Composition.

**Figure 2:**
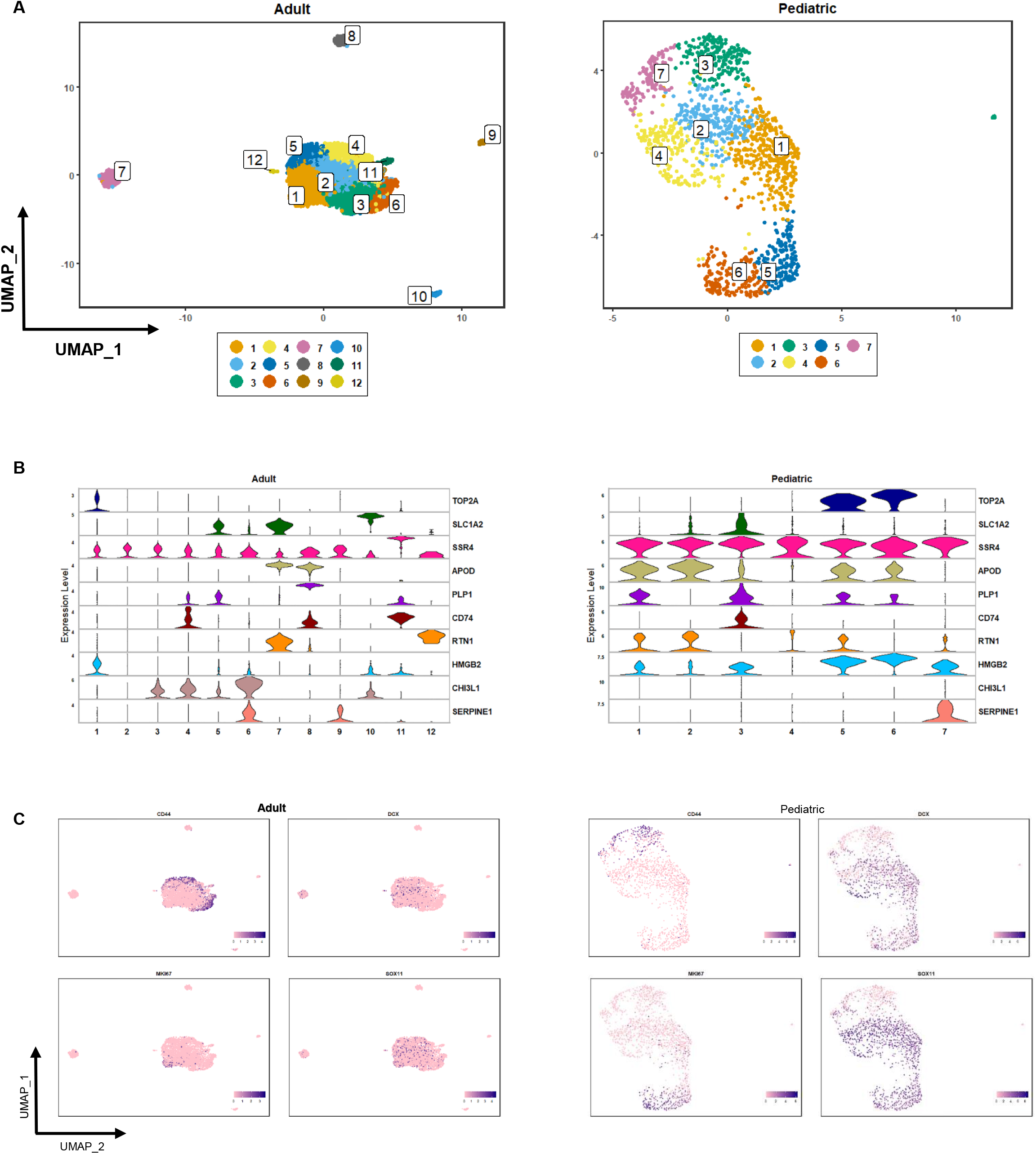
Cell type comparison of Adult and pediatric groups (A) UMAP representation of Adult and Pediatric groups. Clusters are numbered and color coded. (B) Violin plot of cluster wise expression distribution of cell cycle and neural cell type markers. (C) UMAP representation of expression distribution of markers for quiescence, proliferation and migration in adult and pediatric clusters. Color transition represents expression levels with high expression represented in deep blue. All expression values are log transformed.

### Identification of types and cell cycle stages

To determine the presence of cycling and Quiescent cells in both groups, we sought to identify and distinguish these cells from mature neural and glial cells. We used single cell datasets of adult and embryonic brain, from Darmanis, Spyros, et al.[25] as reference dataset and performed unbiased cell type recognition using SingleR package (see methods). The parameters used are described in the methods section. The results confirmed the initial clustering based predictions and showed the presence of both cycling stem cell like (cGSC) and Quiescent stem cell like (qGSC) cells in patient samples from both groups (Figure 3 A). Comparatively, the proportion of cGSCs and qGSCs was much higher in the pediatric group. Specifically, qGSC population was markedly low in adult group. Patient samples BT-S2 and BT-S4 had no identified qGSCs, while only one cell could be identified as qGSC in PJ-32, while PJ-35 had highest number of both qGSCs and cGSCs in the adult group (Figure 3 D), probably because a large number of cells in the group are from PJ-35. Expectedly, both these populations were absent in the non-GBM metastatic tumor sample (GS-15), confirming their importance as key originators of GBM.

**Figure 3:**
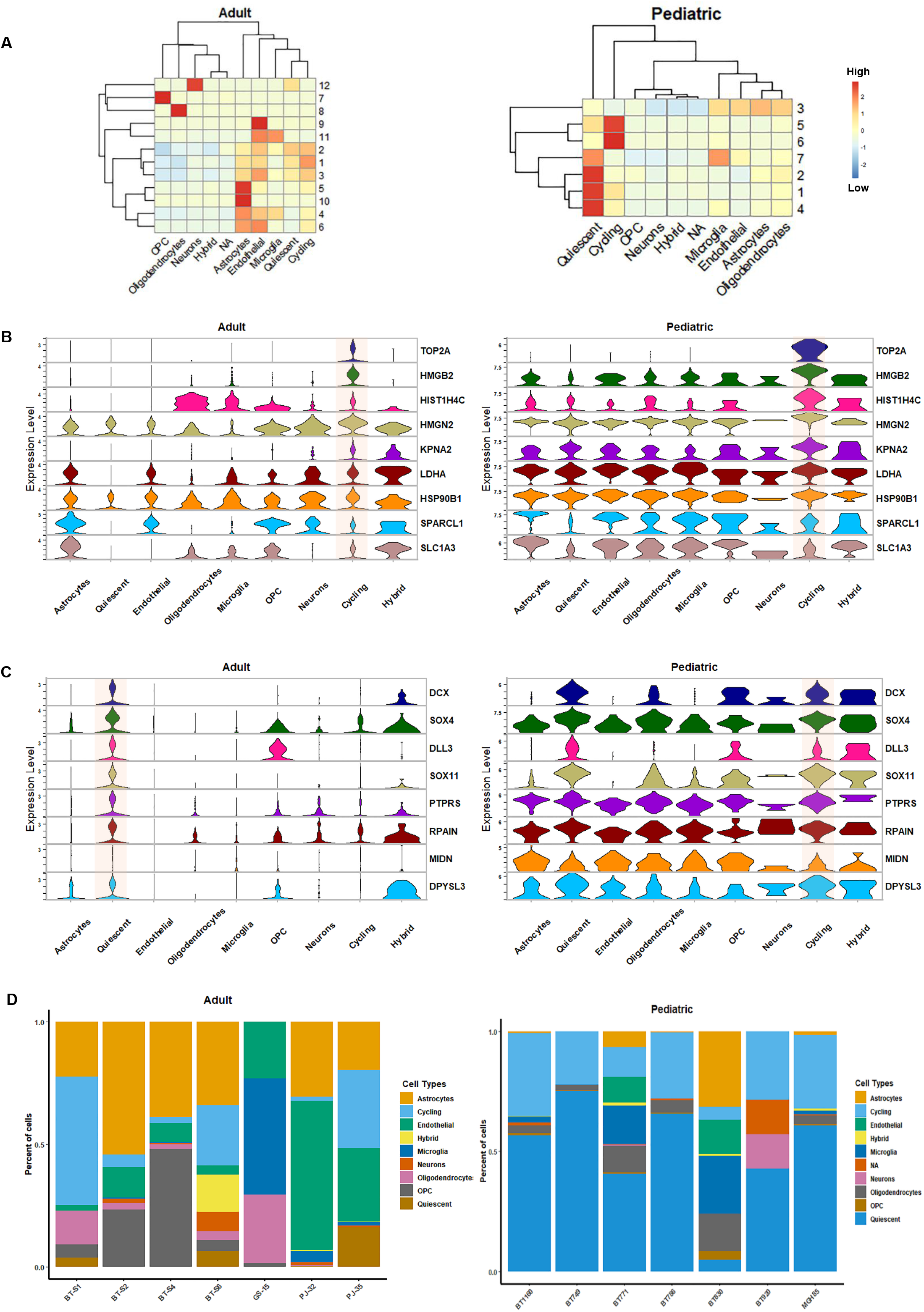
Group wise comparison of identified cell types (A) Heat map of cluster wise expression comparison of markers for cell types in adult and pediatric groups. Cells with no strong matches are marked as NA. (B) Violin plot of top nine overexpressed genes in the cycling cells common to both adult and pediatric groups. (C) Violin plot of top eight overexpressed genes in the quiescent cells common to both adult and pediatric groups. (D) Patient sample wise composition of cell types in adult and pediatric groups. All expression values are log transformed.

### Analysis of cycling and quiescent cells

To do a comparative analysis of the gene expression patterns between the groups, differential gene expression analysis was performed using Model-based Analysis of Single Cell Transcriptomics (MAST) package from R. Significant differentially expressed genes for both qGSCs and cGSCs were compared with other cell types in both adult and pediatric groups (figure 3 B). In both groups, qGSCs had a markedly high expression of *DCX, SOX4, SOX11* and *DLL3* genes. While SOX4 and SOX 11 are both critical in the development and maintenance of neural pluripotent cells, DCX is an essential factor in neurogenesis in neuronal migration. DLL3 is a ligand for the Notch pathway and plays a pleotropic role in notch pathway regulation [32]. On the other hand, cGSCs in both groups were marked by overexpression of HMGB2, HSP90B1 and KPNA2 apart from TOP2A (figure 3 C). HMGB2 is a member of the high mobility protein family, functioning as a modulator of chromatin structure. However, recent study has shown its role in transition from quiescence to activated state in neuronal stem cells (NSCs) [27]. Similarly, HSP90B1, a member of the heat shock protein family, has a role in maintaining embryonic pluripotency [36], whereas KPNA2 is known to be associated with a number of cancers[37]. comparatively, in the adult group, qGSCs and cGSCs have a marked difference in their expression profiles, but less so in the pediatric group. A possible reason for this distinction perhaps is the fact that pediatric brain cells are primed for development.

This is also evident from the cell cycle stage prediction. Previous studies have shown that pluripotency in stem cells is intricately related with cell cycle stages. Whilst a short or truncated G1 (gap1) phase is considered a hallmark of pluripotent state, lengthening of G1 phase is observed when the cells enter cycling phase of rapid differentiation[33–35]. To further confirm the cellular states of these populations, we did a cell cycle state pseudotime prediction using Tricycle R package as described in the methods section. As, the Quiescent or G0 state is not exclusively defined in continuous cell state pseudotime embedding, we expected to find the qGSC cells to be predicted in the G0/G1 phase range, whilst cGSCs to be in G2/M state range. The results were as expected with qGSC almost exclusively in G0/early G1 state whilst cGSCs in late G1 to M states (Supplementary Figure S1). Interestingly, the distribution of cells within cellular states was continuous showing the presence of cells in intermediate states, indicating a transition between qGSC and cGSC states. As, the stem cell like nature of these clusters was supported by both cell type and cell state analysis, we separated these clusters from other cell types and did a comparative analysis of underlying gene expression patterns between adult and pediatric groups to identify universal expression signatures of qGSCs and cGSCs.

### Cluster analysis of cycling and quiescent cells

qGSCs and cGSCs from both groups were reclustered using the same unbiased approach of batch correction as described earlier. Five clusters were observed in both groups (Figure 4 A). Cell cycle pseudotime analysis of the clusters revealed a clear distinction in the cell cycle phase of the clusters in both groups (Figure 4 B). Differential gene expression analysis of the clusters not only revealed the genes involved in quiescence and activation in both groups (Supplementary Figure S2), but also showed a marked similarity between the clusters. we found that a number of highly overexpressed genes in clusters 1 and 2 from the adult group were also highly expressed in clusters 2 and 3 in the pediatric group (Figure 4 C). while the set of clusters overexpressing a large number of ribosomal genes (RP), especially *RPL23, RPL34, RPS3, RPS13, RPS29* also correlate with qGSC cells, the cGSC dominant set was marked by the overexpression of a number of high mobility group (HMG) genes including *HMGB1* and *HMGB2* and heterogeneous nuclear ribonucleoprotein (hnRNP) genes, notably *HNRNPA3* and *HNRNPD*. Recent research points to intricate relation between ribosomal activity and quiescence in stem cells[38, 39]. Indeed, high level of ribosomal presence can block stem cell differentiation. On the other hand, while HMG and TOP2A transcription regulators represent dynamic cell division, hnRNPs are key factors in pre-mRNA processing and transport. Indeed, based on gene expression pattern and cell cycle pseudotime analysis, a picture of sequential progression between the clusters is indicated with the overexpression of ribosomal proteins positively correlating with true quiescence while the overexpression of HMG and hnRNPs indicating progression into cycling phase. In terms of disease model, it is likely that the mechanism of transition from quiescent to cycling states in GBMs remains similar to that of NSCs. In terms of patient samples, all patient samples from the adult group had the presence of ribosome overexpressing cluster (cluster 1), whereas in the pediatric group the ribosome overexpressing cluster (cluster 2) was absent in BT-830. Similarly, the hnRNP/ HMG overexpressing cluster in the adult group (cluster 2) was absent in PJ-32 (Figure 4 D). This may be because the number of cells included in the study may not represent the total tumoral heterogeneity or that the qGSC and cGSC states are interconvertible. Cluster wise differentially expressed genes for both adult and pediatric groups are included in additional file 2.

**Figure 4:**
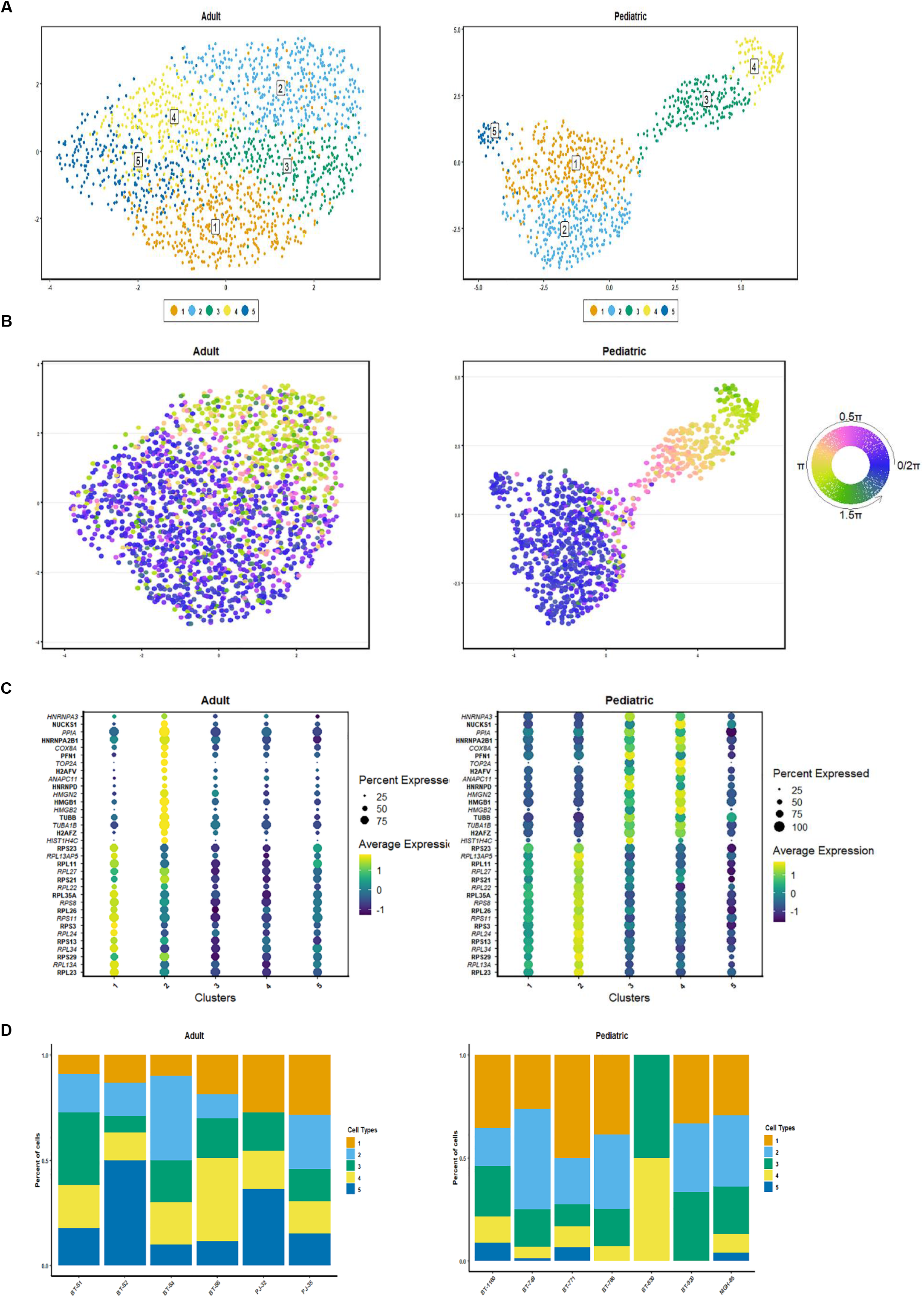
Group wise comparison of cycling and quiescent clusters (A) UMAP representation of integrated clustering of cycling and quiescent cells in Adult and Pediatric groups. Clusters are numbered and color coded. (B) UMAP representation of cell cycle status of the clusters from A. Theta values correspond cell cycle stages as follows: 0.25-1.75π ∼ G0/G1 stage, 0.5π ∼ start of S stage, π ∼ start of G2M stage and 1.5π ∼ middle of M stage. (C) Cluster wise composition of cycling quiescent populations in patient samples from adult and pediatric groups.

Gene ontology (GO) analysis of the clusters further confirmed our observations with the ribosome overexpressing cluster enriched in the biological process of cotranslational protein targeting to membrane or endoplasmic reticulum. The HMG and hnRNPs overexpressing clusters were enriched in cell cycle stages of DNA replication and sister chromatid separation. These clusters are likely representative of cycling cells from S to G2 phases. Interestingly, in the pediatric group, we found that the cluster 5 which comprised of a few cells from samples BT-1160, BT-749, BT-771 and MGH-85 was enriched for neuronal development and differentiation (Figure 5 A). However, we could not determine if this subpopulation is a transformed NSC precursor of a specific lineage.

**Figure 5:**
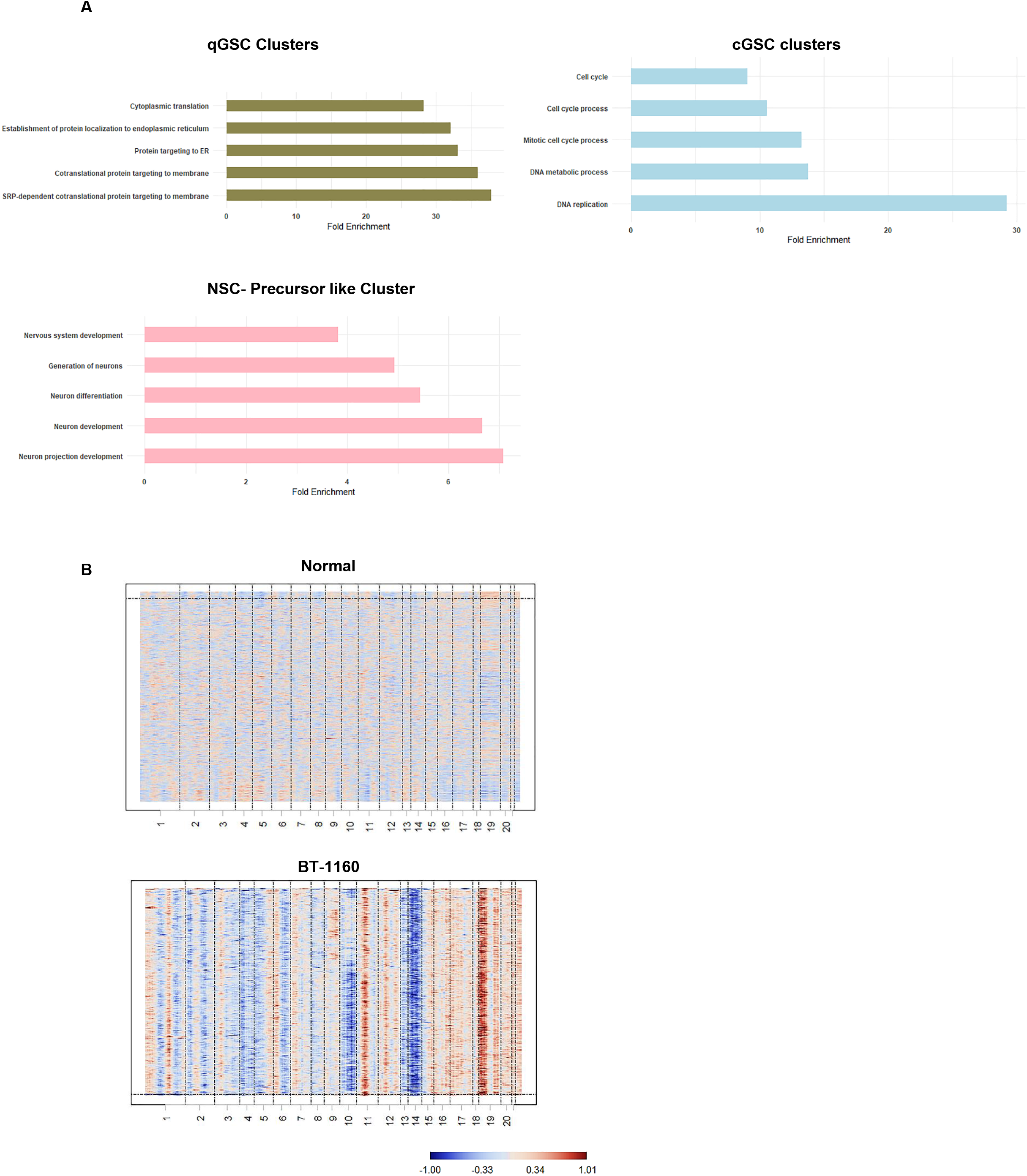
Pathway enrichment and copy number variations in cycling and quiescent cells (A) Bar plots of top five biological processes enriched in cycling and quiescent clusters based on common overexpressed genes in adult and pediatric groups. log fold enrichment of biological processes is represented as bars. (B) Representative images of inferred CNVs in cycling and quiescent cells from one pediatric patient sample (BT-1160) compared to normal cells.

### Analysis of copy number variations (CNVs)

Copy number alterations (gain and/or loss) of the DNA are known to be associated with disease progression in various cancers[40]. Based on the gene expression patterns in the qGSCs and cGSCs, we reasoned that the variations in the chromosomal regions of these cells is likely distinct from normal or differentiated neoplastic glial/neuronal cells. To confirm this hypothesis, we compared the CNV status of GSC (qGSC and cGSC) clusters with non-malignant and differentiated neoplastic cells. For comparison, we used the gene expression counts of 332 of adult normal brain cells from dataset: GSE67835 as reference[25]. The results did not show a clear similarity in CNV patterns of adult and pediatric groups, however, we did observe a pattern of copy number gain at chromosomes 19 and 11 in some patients in the pediatric group (BT-1160, BT-749, BT-771, MGH-85). In terms of pediatric group, we found a consistent CNV pattern in the GSC population (Figure 5 B) (see also supplementary Figure S3). Chromosomes 19 showed a copy number gain while chromosome10 showed a loss of copy number. Locus gain at chromosome 19 is relevant in this study’s context because Chromosome 19 which has a high gene density, also harbors a large number of ribosomal genes [41]. While there seems to be a correlation between copy number alteration at chromosome 19 and ribosomal protein abundance, we could not verify this correlation in terms of causation. However, we consider this an interesting finding, which needs to be explored in detail in the future.

## DISCUSSION

Glioblastomas present marked inter and intra tumoral heterogeneity which is a key hurdle in identifying tumorigenic cell populations and therefore designing robust therapeutic strategies to target them. Recent advances in single cell techniques has helped immensely in studying and intratumoral cellular niches. However, identification of GSCs which are considered to be the drivers of tumor progression and therapy resistance largely remains a challenge because they exist as a small population of cycling and quiescent cells within these niches. The dynamic nature of tumor microenvironment means that these cells show considerable phenotypic plasticity. This behavior of GSCs would suggest that a marker based strategy, although very informative in describing cellular state at a given time, is insufficient in identifying stem cell like tumor populations with marked cellular plasticity. The existence of GSC-like cells in proliferative and quiescent states within these niches is largely agreed upon, however, the identification of quiescent GSC-like population has remained a challenge. Through our study of scRNAseq data from a diverse panel GBM samples we have identified gene signature patterns uniquely associated with cycling and quiescent states in GBM cells.

A number of single cell sequencing based studies in recent years have focused on identifying GSCs[15, 18, 20, 22, 42], these studies support the theory that malignancy in glioblastoma is a function of a small number of progenitor like cells which may exist in neural, mesenchymal, oligodendrocyte or astrocyte like states. The results of this analysis show that although most of the cycling and quiescent GSCs do share similarities with endothelial and glial like cells, they have unique transcriptomic profiles which suggest that these cells are maintained in their own niches within the microenvironment surrounded or populated with mature endothelial, neural and/or glial cells. It is important to underline the observation that while transformed cells from the non–GBM tumor (GS-15) had endothelial, microglia and oligodendrocyte like populations, they lacked both cGSC and qGSC like cells. This provides evidence that glioblastoma is inherently a disease of neural stem/progenitor cells.

A number of studies on cancer stem cells have largely focused on the expression of genes related to a few key transcription factors (Yamanaka factors), OCT4, SOX2, KLF4, and MYC[43] including NANOG which are well known markers for pluripotency. Similarly, makers for proliferation like MKI67 and CD44 are the focus of most of the research on proliferating and pluripotent malignant cells in GBM. However, such approaches overlook the molecular processes involved in maintaining quiescence or triggering proliferation. Indeed, we observed no correlation between CD44 expression and cGSCs. On the other hand, our analysis gives further evidence that SOX4, SOX11 and DCX do overexpress in cycling and quiescent GSCs, however, studying these markers alone won’t explain the molecular process involved in maintaining quiescence or triggering proliferation.

The results of this analysis provides strong evidence that quiescent and cycling stem like cells in GBM share common molecular pathway to maintain quiescence. By comparing the difference in the gene expression profiles of qGSC and cGSC clusters, we have been able to captures the changes in cellular processes of the cells transitioning from quiescent to cycling state. Notably, the quiescent state is underlined by the overexpression of a number of ribosomal genes while the cycling state is marked by the overexpression of HMG and hnRNPs. This switch from ribosomal gene family to HMG and hnRNP genes suggests triggering of entry into cycling phase is accompanied by profound changes in cell physiology. The presence of migration and neuronal differentiation related genes like *S100B, VIM* and *SPARC* in a quiescent like separate cluster (cluster 5) may show the presence of NSC like or progenitor cells. further research is required to understand their interaction with other clusters. In terms of patient groups, the results show that adult tumor samples had a much lower proportion of qGSCs, this is probably reflective of the differences in developmental state of brain. Importantly, the results suggest that qGSCs and cGSCs of NSC nature are a key feature of the glioblastoma tumor microenvironment. These populations are largely interconvertible but are maintained predominantly by the expression of genes distinct to these states.

The role of ribosomal proteins in stem cell maintenance is an area of active research[44]. Recent research on mouse NSCs has shown low protein synthesis rate as a hallmark of quiescent state[45]. This phenomenon of quiescent NSCs is probably due to reduced activity of the mTOR (mammalian target of rapamycin) kinase which acts as a key bridge linking ribosome biogenesis and protein synthesis to induction of pluripotency, self-renewal and differentiation in adult stem cells[46, 47]. Experimentally, it has been shown that knockdown of 4E-BP1 (an mTOR target) promotes differentiation in mouse NSCs[48], mTOR signaling drops when the cell exits cell cycle, leading to suppression of ribosome synthesis, controlling NSC differentiation[49]. It is thus likely that a number of ribosomal proteins are maintained in the quiescent cells to trigger differentiation, thus ensuring and effective transformation of the stem cell state upon receiving environmental signal[47, 48, 50, 51]. While this is a possible theory for the overexpression of ribosomal genes in quiescent cells, further studies are needed to understand this phenomenon in glioblastoma.

In conclusion, this study provides vital insight in the expression profile of cycling and quiescent like cells in glioblastoma. Therapy designs targeting these cells holds great promise in the treatment of GBM patients because studies have shown that these cells are key to developing therapy resistance, migration and proliferation. Targeting quiescent GSCs is critical to overcome tumor relapse. This work is an important step in understanding the molecular processes that govern the quiescent and cycling states in GBMs.

## MATERIALS AND METHODS

### Data resource and selection

All Single-Cell RNA-Seq raw read count matrices and metadata files (wherever available) were were downloaded from Gene Expression Omnibus (GEO) repository. Specifically, gene/cell expression counts from datasets GSE84465[17],GSE117891[18],GSE131928[52] and GSE103224[19] downloaded. The expression matrix for patients included in the study was then curated from the raw expression files. Raw counts for patient data from GSE131928 was not made available by the authors. Log2 transformed count (available) was used instead.

### Data filtration and normalization

All datasets were filtered and analyzed using Seurat V4[53]. Raw data matrix was first filtered using the slandered Seurat protocol to remove possible low quality cells, cells with <200 or >3000 transcripts were excluded from the analysis. In addition, cells of poor quality, recognized as cells with >5% of their transcripts coming from mitochondrial genes, were excluded from the downstream analysis.

### Clustering techniques

Primary clustering of the datasets was done following Seurat protocol. Briefly, after filtering and removal of mitochondrial counts, the data was log normalized and highly variable features were calculated (for this study, we kept the nfeature setting to 6000). Next, the data was scaled before performing linear dimensional reduction. High variable principal components were selected based on percentage variance. Next, the K nearest neighbor graph was constructed based on calculated principal components and clustered. Finally, the dimensional reduction and visualization was done using Uniform Manifold Approximation and Projection (UMAP).

### Differential gene expression analysis

Analysis of differentially expressed genes for each clusters was done by implementing Model-based Analysis of Single Cell Transcriptomics (MAST) package with Seurat[54].

### Integration of datasets

For integrated analysis, following initial Seurat protocol, the fast mutual nearest neighbors (fastMNN) R/Bioconductor package was applied to correct for differences between data sets (batch effect correction)[55]. Clustering was done using default parameters.

### Reference based cell type identification

Cell type identification was performed using singleR[56]. which is an R/Bioconductor package to perform unbiased cell type recognition from single-cell RNA sequencing data, by using reference datasets of pure cell types to identify the cell type of individual single cells independently. Here we used the dataset from Darmanis, Spyros, et al.[25] as reference dataset for cell type identification. Cells not recognized as either of the cell types (NAs) were removed from further analysis.

### Cell cycle trajectory inference

Cell cycle phase of the integrated datasets was inferred using the Tricycle R/Bioconductor package, which uses a fixed reference dataset to infer cell cycle phase of the test dataset[57]. Here, we used the reference dataset provided in tricycle with default parameters to infer cell cycle positions of cells in integrated data. The inferred positions were then project on to the UMAP for visualization. The estimated cell cycle position is bound between 0 and 2pi. The cycle positions approximately relate to theta as: 0.25pi-1.75pi to G0/G1 stage, 0.5pi to start of S stage, pi to start of G2M stage and 1.5pi the middle of M stage.

### CNV analysis

To compare the copy number variations between clusters and datasets, we used CONICSmat (Copy-Number Analysis in Single-Cell RNA-Sequencing from an expression matrix) R package which compares average gene expression of genes within a region to calculate the variance in copy number across samples[58]. Although reference data is not explicitly required, yet, for added certainty, we used normal brain expression matrix of 322 normal brain cells from Darmanis, Spyros, et al[25]. Analysis was done following the default protocol.

### Gene Ontology (GO) analysis

Up to 50 (wherever possible) overexpressing genes for each cluster were analyzed for the enrichment of associated GO terms. Top 5 terms were selected based on fold change and represented graphically.

### Statistical analysis

All data analysis was performed with R. Specific packages used are mentioned in the above sections.

## LIST OF ABBRIVIATIONS

GBM: Glioblastoma
GSC: Glioblastoma stem cell
NSC: Neural stem cell
UMAP: Uniform Manifold Approximation and Projection
cGSC: Cycling glioblastoma stem cell
qGSC: Quiescent glioblastoma stem cell
HMG: High mobility group protein
hnRNP: Heterogeneous nuclear ribonucleoprotein

## ADDITIONAL FILES AND SUPPLEMENTARY FIGURES

Additional File 1: Group wise list of differentially expressed genes according to respective clusters for Figure 2 A.

Additional File 2: Group wise list of differentially expressed genes and common genes according to respective clusters for Figure 4.

Supplementary figure S1: Cell Cycle pseudotime representation of unclustered cycling and quiescent GSCs from adult and pediatric groups.

Supplementary figure S2: Dot plot representation of top ten genes per cluster for figure 4.

Supplementary figure S: CNV profiles of GSCs from pediatric group patient samples.

## Notes

### Competing Interest Statement

The authors have declared no competing interest.

