## Supplementary Figures for "Analysis of single-cell RNA-sequencing data to identify quiescent and proliferating neural cell populations in Glioblastoma"

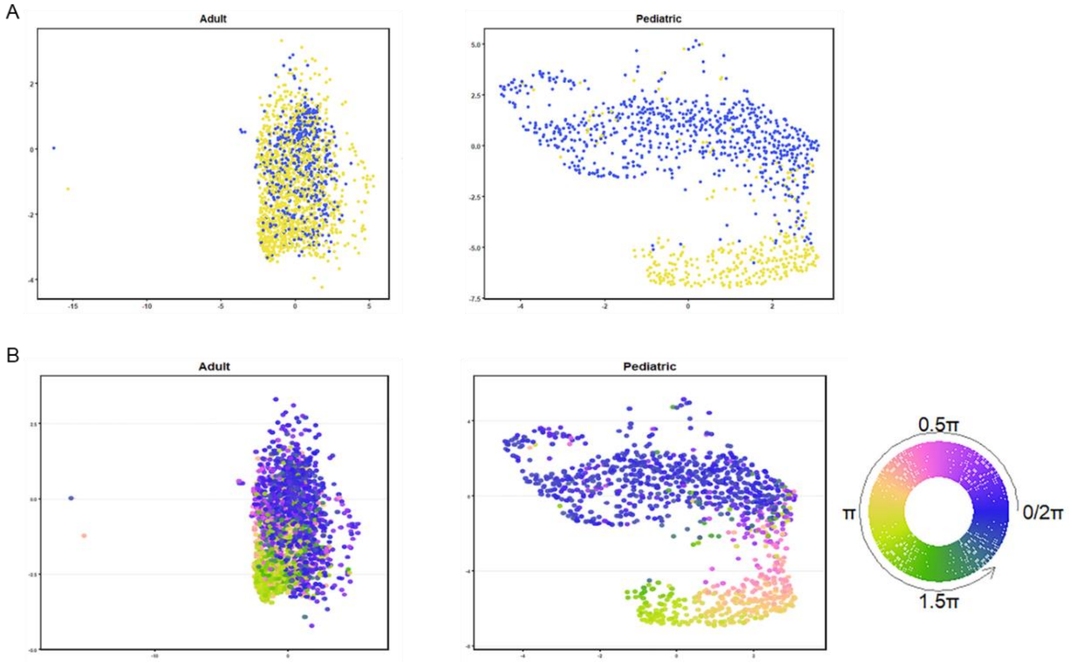


**Figure S1** : (A) UMAP representation of cycling and quiescent cells extracted from adult and pediatric clusters based on cell type prediction. (B) UMAP projection of the same after cell cycle stage prediction showing strong match between both predictions.

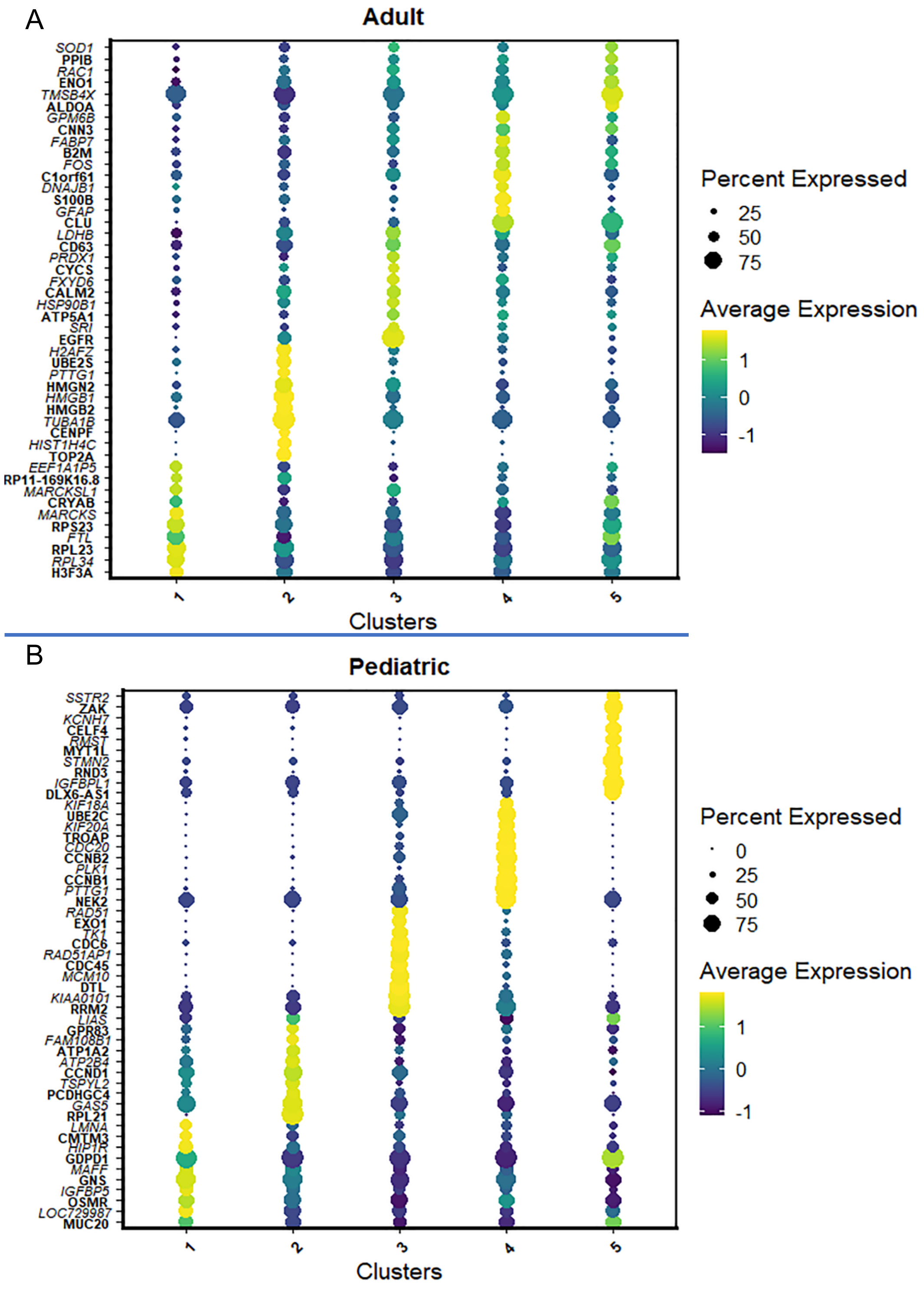


**Figure S2**: Dot plots of gene expression patterns of top ten genes per cluster for adult (A) and pediatric (B) groups. Dot size represents percentage expression; dot color represents average expression (deep blue – low to yellow-high).

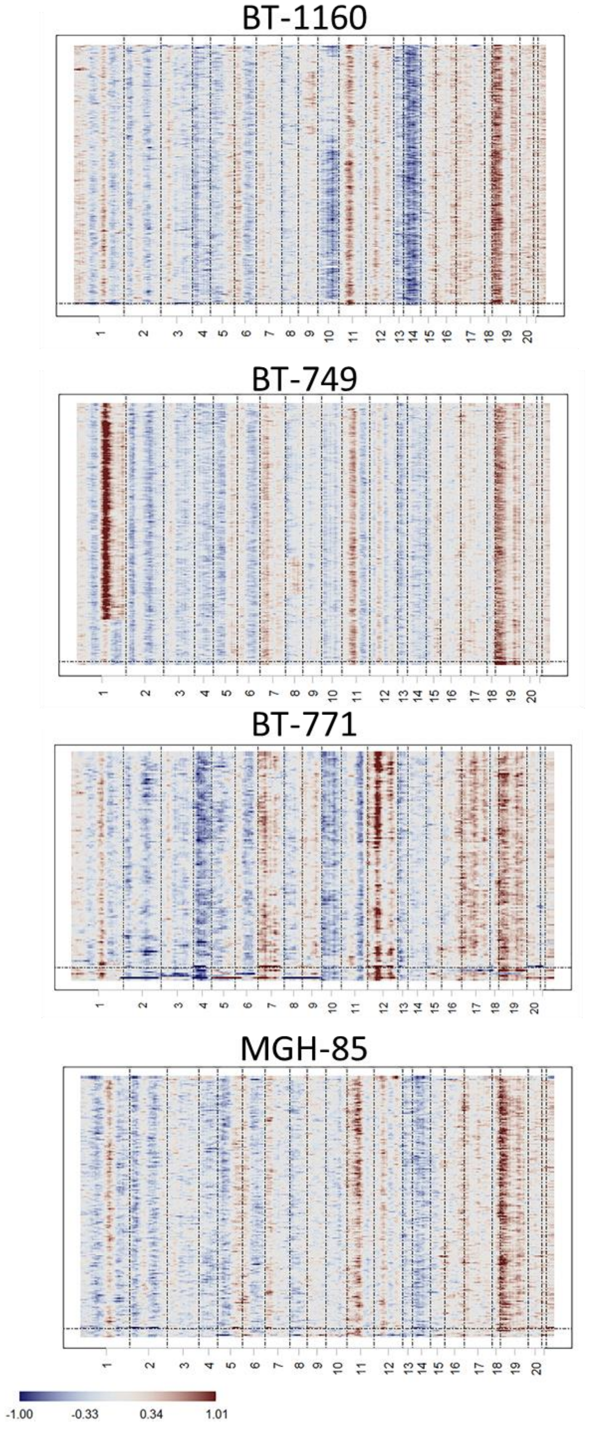


**Figure S3:** CNV profiles of GSCs (qGSC and cGSC) from pediatric patient samples showing a clear pattern of copy number gain at chromosome 19.Color code : Red –gain, Blue –loss.
